# Shifting roles of incomplete lineage sorting and gene flow across the diversification of Old World deer

**DOI:** 10.64898/2026.11.24.768796

**Authors:** Bo Xiao, Xueli Shen, Junxia Yuan, Guilian Sheng

## Abstract

Rapid radiations often generate pervasive phylogenetic discordance through incomplete lineage sorting (ILS) and gene flow, making evolutionary relationships difficult to resolve. Old World deer (subfamily Cervinae) exemplify this process, in which morphological convergence and cytonuclear discordance have long complicated intergeneric relationships and reticulate histories. Here, we analyzed whole-genome resequencing data from 10 individuals representing 10 species across all six extant Cervinae genera. Integrating phylogenomic reconstruction, molecular dating, and genome-wide discordance and allele-sharing tests, we investigated the roles of ILS and introgression. Our nuclear species tree shows that *Axis porcinus* and *Dama dama* diverged sequentially at the base of Cervinae (at 7.69 Ma and 6.56 Ma, respectively), followed by the split between *Cervus* and the *Elaphurus–Rusa–Rucervus* clade at 5.29 Ma, and rapid diversification within *Cervus* during the Early Pleistocene (2.23–1.01 Ma). Discordance patterns shifted across timescales: Miocene divergences were primarily associated with ancestral ILS, whereas Pliocene and Pleistocene divergences showed stronger signatures of gene flow. These included ancient introgression between *Elaphurus davidianus* and ancestral *Cervus*, as well as recent hybridization between *Cervus albirostris* and *Cervus canadensis*. Around the Miocene–Pliocene transition (∼5.3–5 Ma), warmer, wetter conditions promoted woodland recovery, followed by Pleistocene climatic oscillations. These environmental changes likely reshaped habitat connectivity, facilitating secondary contact and gene flow among diverged lineages, particularly during recent radiations. Together, these findings highlight the roles of ancestral polymorphism and episodic gene flow in shaping the reticulate evolutionary history of Cervinae.

## 1. Introduction

Rapid radiation plays a central role in generating extant biodiversity, yet reconstructing the phylogenetic histories of lineages that diverged over brief intervals remains a major challenge (Simões et al., 2016, Whitfield and Lockhart, 2007, Degnan and Rosenberg, 2009). When speciation events occur in close succession, ancestral genetic polymorphisms can persist across multiple splits, resulting in incomplete lineage sorting (ILS) and discordance between gene trees and species trees (Scornavacca and Galtier, 2016, Pamilo and Nei, 1988, Degnan and Rosenberg, 2009). Concurrently, interspecific gene flow, either during ongoing divergence or via subsequent secondary contact, can introduce further genealogical conflict (Hibbins and Hahn, 2022, Mallet et al., 2016). Recent genomic studies have demonstrated that ILS and introgression can jointly shape genomic relationships in rapidly diversifying clades (Guo et al., 2026, Huang et al., 2025, Myers et al., 2024, Rivas-Gonzalez et al., 2023, Feng et al., 2022). However, the relative contributions of these two processes to phylogenetic discordance can vary markedly among taxa with distinct biogeographic backgrounds and temporal scales. Extant Old World deer (subfamily Cervinae) are widely distributed across the Eurasia, spanning latitudes from subtropical to subarctic regions (Pitra et al., 2004). Over the past ∼10 million years, their history of sequential cladogenesis and geographic expansion across contrasting climatic regimes provides an ideal system to examine how ILS and gene flow interact to shape phylogenetic patterns during a continental radiation.

Following the restricted definition of Cervinae (corresponding to the tribe Cervini and excluding muntjacs, Muntiacinae), this subfamily comprises six extant genera: *Axis*, *Dama*, *Cervus*, *Elaphurus*, *Rusa*, and *Rucervus* (Pitra et al., 2004, Groves and Grubb, 1987, Randi et al., 2001). Their classification has long been debated because early morphological studies based on antler structure, dentition, and cranial features often yielded conflicting generic boundaries (Brooke, 1878, Groves and Grubb, 1987, Heckeberg, 2020). Later molecular studies based on mitochondrial DNA and a few nuclear genes prompted major taxonomic revisions: for example, the formerly monotypic genus *Przewalskium* was incorporated into *Cervus* as the white-lipped deer (*Cervus albirostris*), and the red deer complex (*Cervus elaphus* sensu lato) was split into distinct species, including European red deer (*Cervus elaphus*), wapiti (*Cervus canadensis*), and Central Asian red deer (*Cervus hanglu*) (Pitra et al., 2004, Hu et al., 2019). Nevertheless, while *Axis* and *Dama* are generally recognized as distinct, early diverging genera (Pitra et al., 2004, Heckeberg, 2020), the relationships and taxonomic boundaries among *Cervus*, *Elaphurus*, *Rusa*, and *Rucervus* remain unresolved; the latter three have been variably treated as separate genera or subsumed as subgenera within a broadly defined *Cervus* (Heckeberg, 2020, Hassanin et al., 2012).

Beyond these taxonomic uncertainties, pervasive discordance between nuclear and mitochondrial phylogenies complicates Cervinae systematics and suggests recurrent hybridization. Within *Cervus*, nuclear markers generally group *C. elaphus*, *C. canadensis*, and sika deer (*Cervus nippon*) apart from *C. albirostris*, whereas mitochondrial phylogenies place *C. albirostris* and *C. nippon* within the *C. canadensis* clade (Gilbert et al., 2006, Pitra et al., 2004, Hassanin et al., 2012, Heckeberg, 2020). Documented hybridization between *C. nippon* and *C. elaphus* further indicates the potential for interspecific gene flow (Senn et al., 2010, Hu et al., 2019). Another notable case is Père David’s deer (*Elaphurus davidianus*), whose mitochondrial affinity to Eld’s deer (*Rucervus eldii*) contrasts with nuclear, morphological, and karyotypic evidence linking it to *Cervus*. This discordance has led to the hypothesis of an ancient hybrid origin involving ancestral *Rucervus* and *Cervus* lineages (Meijaard and Groves, 2004). These conflicting phylogenetic signals highlight the need to disentangle the contributions of ILS and introgression to the evolutionary history of Cervinae.

In this study, we used whole-genome resequencing data from 10 species representing all six extant genera of Cervinae, with Reeves’s muntjac (*Muntiacus reevesi*) as the outgroup, to investigate the evolutionary history of this subfamily. We reconstructed species relationships and divergence times using phylogenomic analyses and molecular dating, and combined genome-wide analyses of phylogenetic discordance and allele sharing to investigate the roles of ILS and introgression. Specifically, we aimed to (i) resolve the phylogenetic relationships and divergence history of Cervinae; (ii) assess the contributions of ILS and post-divergence gene flow to phylogenetic discordance within and among genera; and (iii) characterize the genomic patterns of reticulate evolution, with particular attention to the proposed hybrid origin of *Elaphurus davidianus* and the rapid diversification of *Cervus*.

## 2. Materials and Methods

### 2.1. Data collection

We collected raw sequencing reads of 10 genomes representing 10 extant Cervinae species for phylogenomic analyses (Figure 1). The sampled species represented six genera within Cervinae: *Cervus* (*n* = 5), *Rusa* (*n* = 1), *Rucervus* (*n* = 1), *Elaphurus* (*n* = 1), *Dama* (*n* = 1), and *Axis* (*n* = 1). For species with multiple available genomic datasets, we selected individuals with the highest sequencing depth. Most selected genomes had an average sequencing depth greater than 12×, except for the *Rucervus* individual, which had a sequencing depth of approximately 3×. In addition, raw sequencing reads of a *M. reevesi* genome was obtained and used as the outgroup.

**Figure 1.**
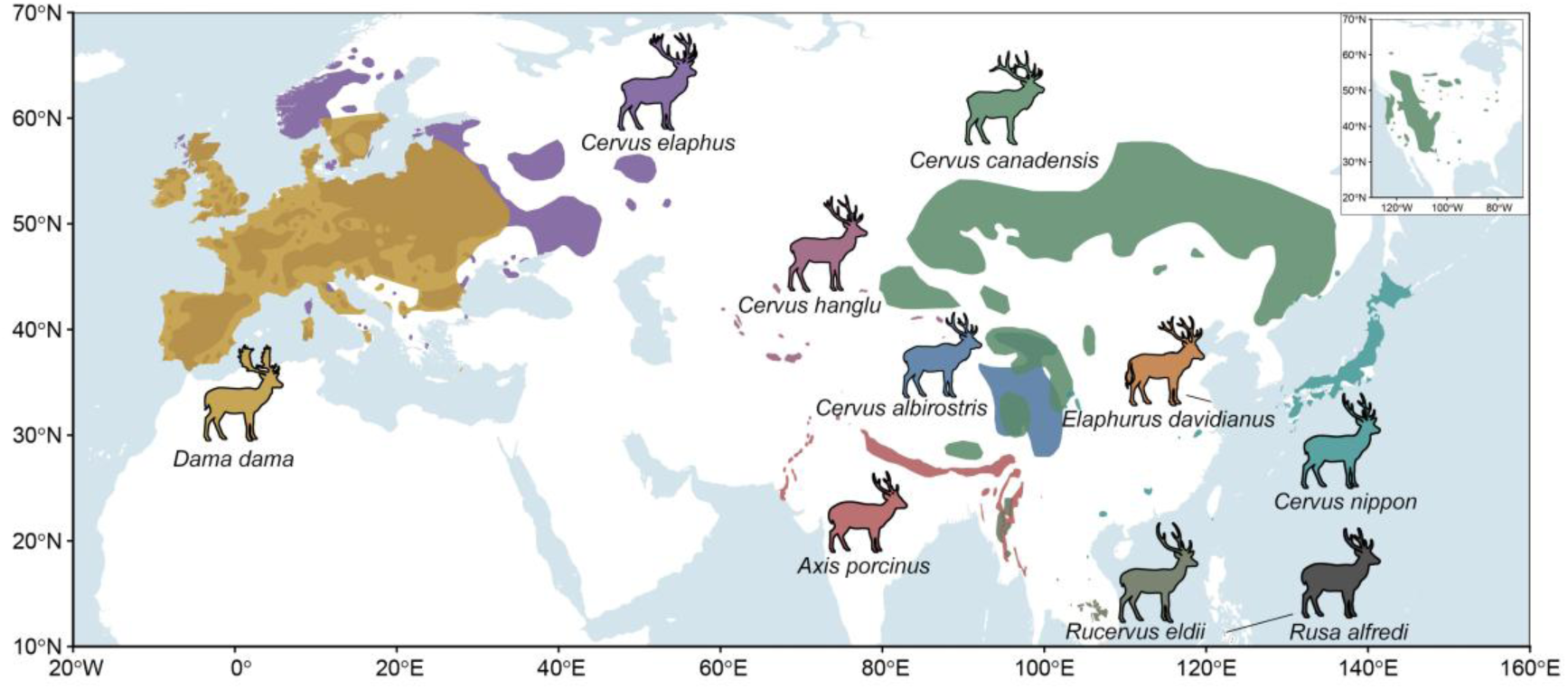
Geographic ranges and silhouette illustrations of the 10 Cervinae species analyzed in this study. Distinct colors represent their contemporary distribution ranges. The inset map in the upper right corner displays the North American range of *Cervus canadensis*. Geographic distribution data were compiled from the IUCN Red List of Threatened Species.

All sequencing raw data were downloaded from publicly available Sequence Read Archive (SRA) repositories, including those hosted by the National Center for Biotechnology Information (NCBI) (Bergeron et al., 2023, de Jong et al., 2025, Chen et al., 2019, Barnard et al., 2023, Mudd et al., 2020) and National Genomics Data Center (NGDC) (Zhong et al., 2025, Zheng et al., 2024). Detailed information on sample sources, accession numbers, sequencing platforms, and genome coverage is provided in Supplementary Table S1.

### 2.2. Raw data processing

Raw sequencing reads were processed using fastp v0.24.0 (Chen et al., 2018) to remove adapter contamination, low-quality bases, and short reads. Adapter trimming and poly-G tail removal were enabled, and reads were retained only when the base quality score was ≥25 and the read length was ≥30 bp after filtering. The resulting clean reads were used for subsequent alignment analyses.

Clean reads were independently aligned to the nuclear genome reference assembly of Chinese water deer (*Hydropotes inermis*) (accession number: GCA_056247825.2) (Wu et al., 2026) and the mitochondrial genome reference (accession number: NC_011821.1) using BWA v0.7.18 (Li and Durbin, 2009) and the mem algorithm, and only mapped reads with a mapping quality score (MAPQ) ≥30 were retained. The resulting BAM files were sorted and processed using GATK v4.0 MarkDuplicates to identify and mark duplicate reads. The final deduplicated BAM files were used for downstream genomic analyses.

For mitochondrial phylogenetic analyses, mitochondrial consensus sequences were generated from the deduplicated BAM files using ANGSD v0.941 (Korneliussen et al., 2014) with the ‘-doFasta 2’ option. Only uniquely mapped reads with mapping quality ≥30 and bases with quality scores ≥30 were considered for consensus sequence reconstruction. Nucleotide sites with sequencing depth lower than 5 were excluded, and allele counts were generated during the consensus sequence generation process.

### 2.3. Mitochondrial phylogenetic analysis

Mitochondrial dataset consisted of 16,355 bp of sequences from 10 Cervinae individuals (representing 10 species) and one outgroup *M. reevesi*. A maximum-likelihood (ML) phylogenetic tree was reconstructed using IQ-TREE v3.0.1 (Wong et al., 2026) with the best-fit nucleotide substitution model (TIM2+F+I+G4) automatically selected by the program. Branch support was assessed using the ultrafast bootstrap method with 1,000 replicates. The resulting phylogenetic tree was visualized and edited using FigTree v1.4.3 (Rambaut et al., 2016).

### 2.4. Single nucleotide polymorphisms (SNPs) calling and filtering

SNP calling was performed using bcftools v1.21 (Danecek et al., 2021). For each chromosome, reads mapped to the reference genome were processed using bcftools mpileup, followed by variant calling with bcftools call –mv. The resulting VCF files were merged across autosomes using bcftools concat. We retained biallelic SNPs with a quality score (QUAL) ≥ 30 using bcftools view, and sites with a missingness rate >20% were excluded. Transposable element (TE) regions were subsequently removed using bedtools intersect –v based on the annotated TE coordinates (Wu et al., 2026).

### 2.5. Species tree reconstruction

To reconstruct the coalescent-based species tree of Cervinae, the TE-masked SNP dataset was first converted into the haplotype format using parseVCF.py (available at https://github.com/simonhmartin/genomics_general). ML gene trees were reconstructed from genomic windows using phyml_sliding_windows.py from the same package. Windows were defined using a size of 50 kb and a step size of 1,000 kb, and only windows containing at least 20 variable sites were retained (*n*=2,323) for phylogenetic reconstruction. PhyML v3.4 (Guindon et al., 2010) was used to infer maximum-likelihood trees, with topology, branch lengths, and substitution model parameters optimized (--optimise tlr). The resulting gene trees were subsequently used to infer the species tree with ASTRAL-III (Zhang et al., 2018).

### 2.6. Divergence time estimation

Consensus sequences for each individual were generated from BAM files using ANGSD with the following parameters: –dofasta 2 –setMinDepth 5 –minmapq 30 –minq 30 –uniqueonly 1 –only_proper_pairs 1 –docounts 1. For *R. eldii*, which had lower sequencing coverage, the minimum depth threshold was reduced to 3 (-setMinDepth 3). We then extracted the corresponding sequences from all valid genomic windows used for species tree reconstruction using the getfasta function in bcftools v1.21. The resulting sequences were aligned across individuals for each window and concatenated into a single alignment using a custom Python script (fasta2phy.py) to generate the input file for MCMCtree.

Divergence times were estimated using MCMCtree implemented in PAML v4.10.10. We employed the approximate likelihood calculation method. Briefly, the likelihood gradient and Hessian matrices were first calculated using usedata = 3 to generate the out.BV file. This file was subsequently used as input for the MCMC analysis with usedata = 2, from which posterior distributions of divergence times were obtained. An independent-rates relaxed molecular clock model (clock = 2) and the GTR nucleotide substitution model (model = 7) were used, with five discrete categories for among-site rate variation under a gamma distribution (alpha = 0.5, ncatG = 5). The birth-death parameters were set to 0.01, 0.01, and 0.1. Gamma priors were specified for the transition/transversion rate ratio (kappa_gamma = 6 2), the gamma shape parameter (alpha_gamma = 1 1), the mean substitution rate (rgene_gamma = 2 2000 1), and the rate-drift parameter (sigma2_gamma = 1 10 1). The divergence between Cervinae and *M. reevesi* was used as the root calibration, with an age constraint of 8.31–11.85 Ma (median = 10.08 Ma). Each MCMC run consisted of 100,000 burn-in iterations, followed by 100,000 samples collected at intervals of 1,000 iterations. Two independent MCMC runs were performed, and their consistency was assessed before combining posterior samples to estimate node ages and 95% credible intervals.

The initial divergence time analysis included all ten sampled Cervinae individuals and one *M. reevesi* outgroup. To evaluate whether the low-coverage *R. eldii* sample affected divergence time estimates, we repeated the analysis after excluding this individual, resulting in a dataset comprising nine Cervinae individuals and the same outgroup. Two independent MCMC runs were also performed for this reduced dataset. For nodes shared between the two analyses, the relative change in divergence time was calculated as the difference between the posterior mean ages from the reduced and full datasets, divided by the posterior mean age from the full dataset and expressed as a percentage. Positive values indicate older divergence-time estimates after excluding *R. eldii*, whereas negative values indicate younger estimates.

### 2.7. PhyTop analysis

PhyTop (Shang et al., 2024) was performed to investigate the distribution of alternative phylogenetic topologies across the genome and assess the potential contributions of ILS and introgression/hybridization (IH) to phylogenetic discordance. The analysis was conducted using the above inferred species tree and genomic window-based phylogenetic trees following the recommended procedure of PhyTop.

### 2.8. The *f*-branch statistics

Potential introgression events among Cervinae species were investigated based on the TE-masked SNP dataset, using Dsuite v0.5 (Malinsky et al., 2020). The D-statistic (ABBA-BABA test) was calculated for all possible combinations of three ingroup taxa using the outgroup species. The resulting D-statistics were further analyzed using the *f*-branch statistic, with the inferred species tree provided as the reference topology, to identify potential introgression signals across the phylogeny.

### 2.9. Sliding-window topology analysis

To investigate phylogenetic discordance across the genome, maximum-likelihood phylogenetic trees were reconstructed using sliding windows of 50, 100, 500, and 1,000 kb. Only gap-free windows containing at least 20 variable sites were retained for tree reconstruction. Maximum-likelihood trees were inferred using PhyML v3.4 (Guindon et al., 2010) with topology, branch lengths, and substitution model parameters optimized (--optimise tlr).

Topology weighting analysis was performed using twisst.py (available at https://github.com/simonhmartin/twisst) to quantify the relative frequencies of alternative phylogenetic topologies for selected quartets of species. The resulting topology weights were further processed using custom R scripts to calculate the relative contributions of different topologies and identify consecutive genomic windows supporting the same topology, thereby characterizing the lengths of genomic segments with consistent phylogenetic signals.

### 2.10. Phylogenetic analysis based on LD-pruned SNPs

To provide an independent assessment of genome-wide genetic relationships among species while reducing the influence of linkage disequilibrium, an additional phylogenetic analysis was performed using LD-pruned SNPs. SNPs were pruned using PLINK v1.9 (Chang et al., 2015) with a sliding window of 50 SNPs, a step size of 5 SNPs, and an (r*^2^*) threshold of 0.2. Pairwise genetic distances were then calculated from the LD-pruned VCF file using VCF2Dis (Xu et al., 2025), and the resulting distance matrix was used to construct a neighbor-joining (NJ) tree.

### 2.11. Genome-wide allele sharing analysis

Genome-wide allele sharing patterns were investigated using ANGSD v0.941 (Korneliussen et al., 2014) based on BAM files. D-statistics were calculated using – doAbbababa 1, with the inferred species tree used to define all possible combinations of H1, H2, and H3 taxa, with the outgroup specified as H4. Only autosomal regions were analyzed. Reads with mapping quality below 30 and bases with base quality below 30 were excluded, and only uniquely mapped reads were retained. A block size of 1 Mb was used for calculating D-statistics. The resulting statistics were used to identify asymmetric allele sharing among different species combinations across the genome.

### 2.12. Genome-wide pairwise genetic distance analysis

Genome-wide pairwise genetic distances among individuals were estimated using ANGSD v0.941 (Korneliussen et al., 2014). The analysis was also restricted to autosomal regions. Only uniquely mapped reads with mapping quality and base quality of at least 30 were retained, and reads flagged as problematic were removed. Genotype counts were calculated using –doCounts 1, and pairwise identity-by-state (IBS) distances were estimated using –doIBS 2 to generate a pairwise distance matrix (− makeMatrix 1). As a minimum sequencing depth of 5 was required at each site, and sites covered in all individuals were retained for analysis, the low-quality individual of *R. eldii* was excluded from this analysis.

## 3. Results

### 3.1. Phylogenetic relationships and divergence time of Cervinae

Based on 118,903,837 autosomal SNPs, the nuclear species tree inferred by ASTRAL from 2,323 window-based ML trees resolved the relationships among the 10 Cervinae species. Time calibration of the nuclear species tree placed the divergence between Cervinae and the outgroup *M. reevesi* at approximately 10.23 Ma during the Miocene (Figure 2A, B). Within Cervinae, the basal lineages hog deer (*Axis porcinus*) and fallow deer (*Dama dama*) diverged sequentially at 7.69 Ma and 6.56 Ma, respectively. The split between *Cervus* and the clade comprising *Elaphurus*, *Rusa*, and *Rucervus* occurred near the Miocene–Pliocene boundary at 5.29 Ma. Within the latter clade, *E. davidianus* diverged first at 4.47 Ma, followed by the split between Visayan spotted deer (*Rusa alfredi*) and *R. eldii* at 3.36 Ma during the Pliocene. Diversification within the genus *Cervus* occurred primarily during the Early Pleistocene, beginning with the divergence of *C. nippon* at 2.23 Ma and *C. albirostris* at 1.56 Ma, followed by the radiation of the red deer group (*C. elaphus* at 1.23 Ma, and *C. hanglu* and *C. canadensis* at 1.01 Ma).

**Figure 2.**
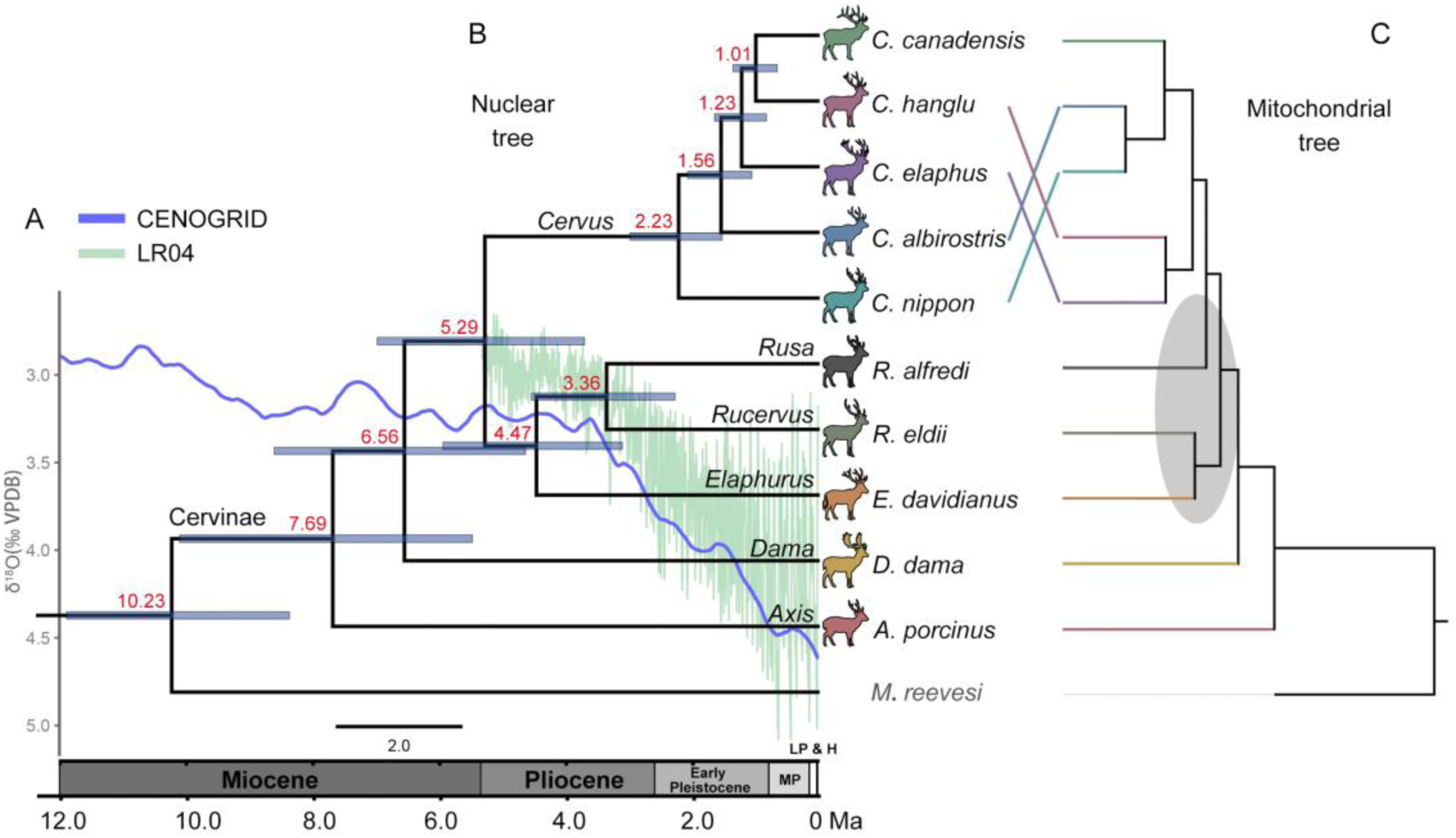
Time-calibrated nuclear phylogeny, paleoclimate curves, and cytonuclear discordance in Cervinae. (A) Global benthic δ^18^O climate records from the CENOGRID (blue) (Westerhold, 2020) and LR04 (green) (Lisiecki and Raymo, 2005) stacks over the past 12 Ma. Geological epochs are shown along the bottom axis: Miocene, Pliocene, Early Pleistocene, Middle Pleistocene (MP), and Late Pleistocene and Holocene (LP & H). (B) Time-calibrated nuclear species tree inferred based on the coalescent species tree topology. Red numbers at internal nodes indicate mean divergence estimates, and horizontal light blue bars represent the 95% highest posterior density (HPD) intervals. Representative silhouettes illustrate morphologies of the 10 Cervinae species. (C) Mitochondrial tree inferred from complete mitochondrial sequences. Within the genus *Cervus* and closely related taxa, marked topological rearrangements are observed among five *Cervus* species as indicated by crossing lines. Shaded ovals highlight significant phylogenetic incongruences. *Muntiacus reevesi* served as the outgroup.

Duplicate divergence time estimates for both the 10-species dataset and the reduced 9-species dataset (excluding *R. eldii*) produced almost identical posterior distributions (Figure S1A). Across all eight shared internal nodes, the 95% HPD intervals overlapped broadly between the two datasets. Relative changes in mean node ages ranged from −6.5% to +0.6%, with seven of the eight nodes differing by −2.7% to +0.6% (Figure S1B). The largest shift (−6.5%) occurred at node 4, which directly subtends the *Elaphurus*–*Rusa*–*Rucervus* clade.

In the nuclear species tree, *R. alfredi*, *R. eldii*, and *E. davidianus* formed a monophyletic group. In contrast, *R. alfredi* was not recovered as part of the same clade as the latter two species in the mitochondrial tree (Figure 2C). Another major incongruence occurred within *Cervus*. In the nuclear species tree, *C. nippon* and *C. albirostris* were both positioned outside the red deer complex. In the mitochondrial tree, however, these two species clustered with *C. canadensis*, forming a clade sister to the group comprising *C. hanglu* and *C. elaphus*.

An additional NJ tree (Figure S2) constructed from 10,041,723 LD-pruned SNPs showed several topological differences from the ASTRAL species tree, particularly in the positions of *E. davidianus* and *C. hanglu*. In the NJ tree, *E. davidianus* was placed at the base of *Cervus* rather than grouping with *R. alfredi* and *R. eldii*. Meanwhile, *C. hanglu* clustered with *C. elaphus* instead of *C. canadensis*, resembling the mitochondrial topology in this respect.

Pairwise autosomal genetic distances among the nine Cervinae species further revealed substantial variation in genetic divergence (Figure S3). The mean pairwise genetic distance across Cervinae was 0.01322. *A. porcinus* and *D. dama*, which occupied basal positions in the species tree, showed greater genetic distances from the other species than the overall mean. By contrast, pairwise distances among the five *Cervus* species were generally lower than the subfamily-wide mean. Notably, the genetic distance between *C. albirostris* and *C. canadensis* was comparable to the genetic distances among the three red deer species.

### 3.2. Genome-wide allele sharing and potential introgression

The *f*-branch analysis revealed pervasive gene flow across the Cervinae phylogeny, with most events showing relatively low values of *f_b_* (< 0.05) (Figure 3). Substantial introgression signals were detected primarily within the genus *Cervus*. Specifically, the highest *f_b_* value occurred between *C. albirostris* and *C. canadensis* (*f_b_* > 0.2), accompanied by clear signals between *C. albirostris* and *C. nippon*, as well as between *C. albirostris* and the ancestral branch of *C. canadensis* and *C. hanglu*. Marked introgression was also observed between *C. nippon* and *C. canadensis*, as well as between *C. hanglu* and *C. elaphus*. Furthermore, *E. davidianus* displayed broadly comparable *f_b_* estimates with different *Cervus* species.

**Figure 3.**
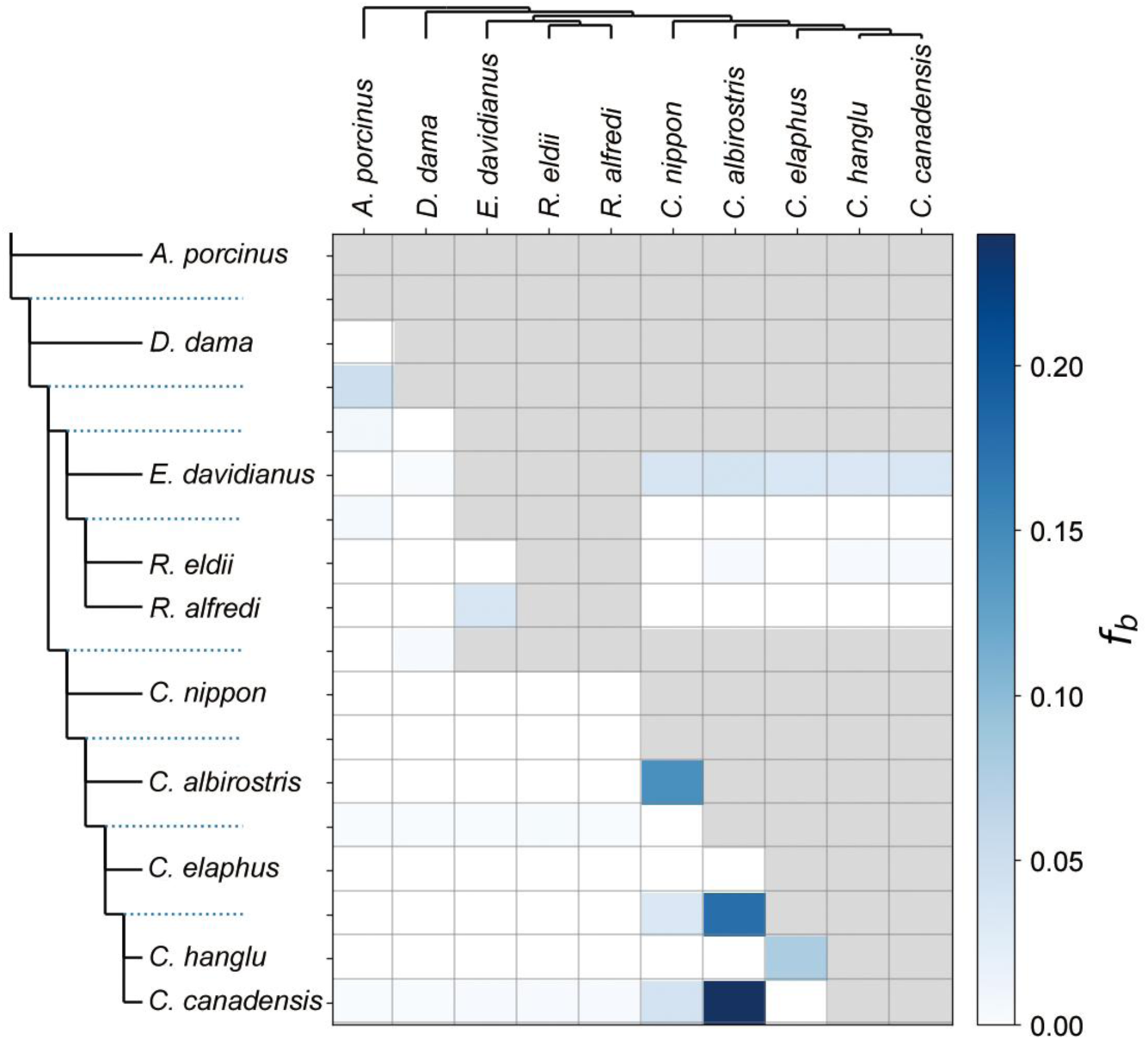
Genome-wide excess gene flow and hybridization among Cervinae lineages calculated by *f*-branch statistics. *f*-branch metrics quantifying gene flow across the Cervinae phylogeny based on nuclear genomic data. The tree at the top represents the recipient taxa, while the tree on the left depicts the donor lineages, with internal nodes mapped to corresponding ancestral branches (dotted lines). Blue tiles indicate significant excess allele sharing (*f_b_* > 0) between donor branches and recipient species, with color intensity proportional to the *f_b_* value. Grey squares denote test combinations that are topologically unassessable due to phylogenetic constraints.

D-statistics calculated using ANGSD yielded congruent patterns (Figure S4). Across all 120 assessed taxon combinations, 113 trios (94%) exhibited significant D values (|Z| ≥ 3). In line with the *f*-branch results, significant excess allele sharing was identified among *C. albirostris*, *C. nippon*, and *C. canadensis* within *Cervus* (Figure S5), with D values ranging from 0.0496 to 0.3266. In tests evaluating gene flow between *E. davidianus* and *Cervus* (Figure S6), all five *Cervus* species shared comparable levels of excess alleles with *E. davidianus*. In trios of ((*R. eldii*, *E. davidianu*s), *Cervus*) and ((*R. alfredi*, *E. davidianus*), *Cervus*), D values ranged from 0.1381 to 0.1428 and from 0.0928 to 0.1148, respectively. By contrast, trios placing *E. davidianus* as H3 against pairwise combinations of *Cervus* species yielded much lower D values (0.0076–0.0682).

### 3.3. Dissecting ILS and IH across Cervinae

PhyTop analysis of local window-tree topologies across internal nodes revealed distinct patterns of ILS and IH across internal nodes of the species tree (Figure 4A, B). Across the evaluated divergences, the frequency distribution of discordant topologies revealed distinct evolutionary dynamics between intergeneric splits and diversification within the genus *Cervus* (Figure 4A).

**Figure 4.**
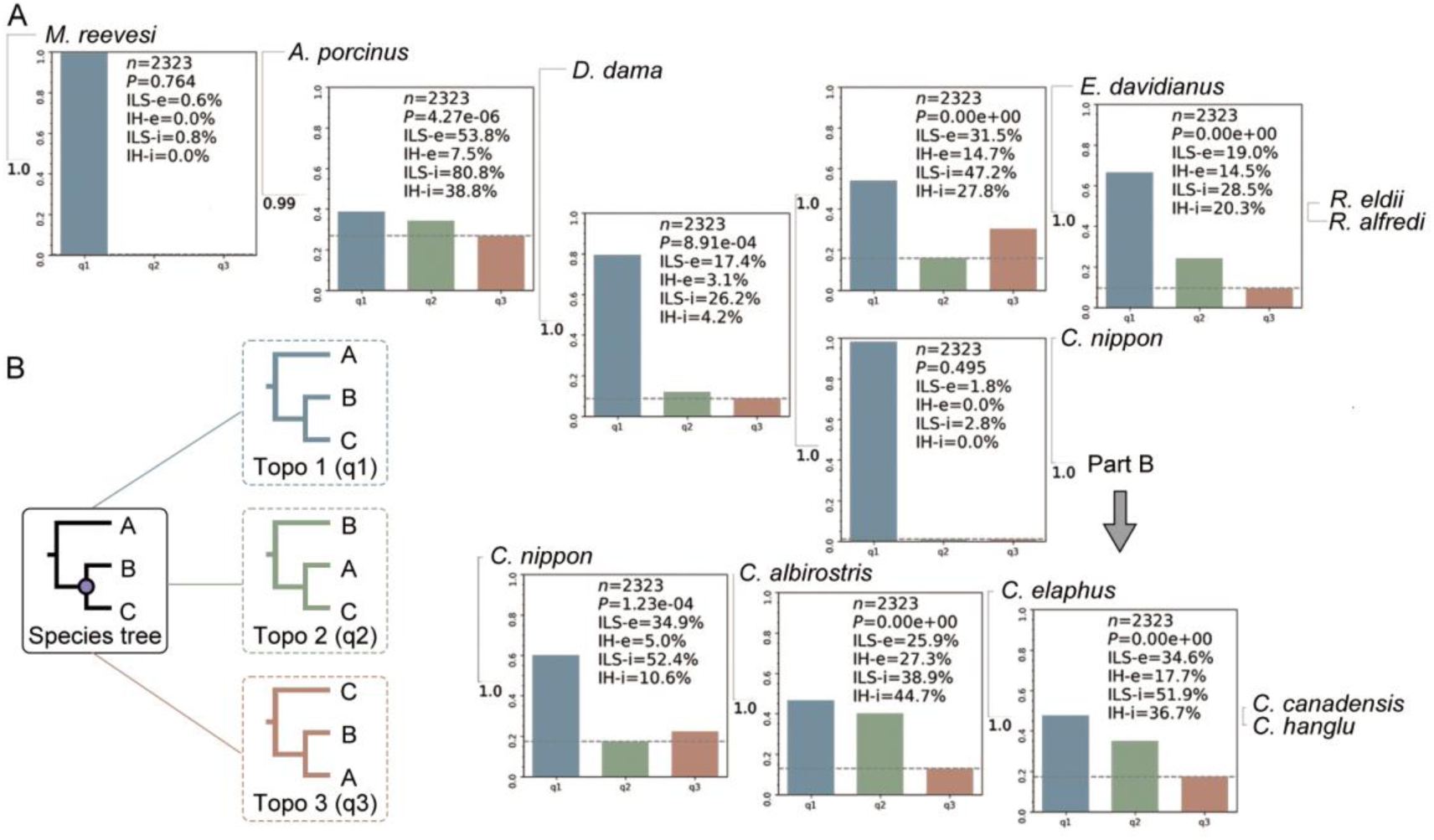
Disentangling ILS and IH across 10 Cervinae lineages inferred by PhyTop. (A) Genome-wide topological distributions and estimated ILS and IH indices across internal nodes of the Cervinae species tree, based on 2,323 sliding-window ML trees (50 kb window size, 1,000 kb step size). Bar charts illustrate the observed frequencies of the concordant species-tree topology (q1) and two discordant alternative topologies (q2 and q3). The grey dashed lines denote the neutral expectation (1/3) under pure ILS. Inset text boxes report tree counts (*n*), *p*-values from *χ*^2^ tests of topological symmetry between q2 and q3, proportions of discordance explained by ILS and IH under equal branch lengths (ILS-e, IH-e), and model-based strength indices under independent branch lengths (ILS-i, IH-i). (B) Schematic illustrations of the three alternative topologies for a rooted three-taxon statement (((A, B), C), Outgroup): the concordant topology (q1: (A, (B, C))) and two discordant topologies: q2: (B, (C, A)); q3: (C, (B, A)).

At deep intergeneric divergences, topological discordance was either negligible or dominated by ILS without asymmetric gene flow. The split of the outgroup *M. reevesi* showed near-complete topological concordance with minimal conflict, representing a well-resolved node (ILS-i = 0.8%, IH-i = 0.0%). For the divergence of *A. porcinus*, although topological conflict was pervasive, the two alternative topologies occurred at comparable frequencies near the neutral expectation (1/3), where discordance was predominantly driven by high levels of ILS rather than IH. Similarly, the node subtending *D. dama* was well resolved, with the species-tree topology accounting for most genomic windows and showing negligible introgression signals (IH-i = 4.2%). By contrast, the node linking *E. davidianus*, *R. eldii*, and *R. alfredi* exhibited significant topological asymmetry (*p* < 0.01), indicating that the observed discordance arose from a combination of ancestral ILS and substantial intergeneric introgression.

Within the genus *Cervus*, topological discordance was pronounced and heavily shaped by gene flow. At the node defining *C. albirostris*, the concordant species-tree topology was recovered in fewer than half of the genomic windows. The alternative topologies displayed severe asymmetry (*p* < 0.001), with one discordant topology occurring almost as frequently as the species-tree topology, leading to a high estimated IH-i value (44.7%) that exceeded the ILS-I (38.9%) (Figure 4A). Marked asymmetry between alternative topologies was likewise detected within the red deer complex (*C. canadensis*, *C. hanglu*, and *C. elaphus*), reflecting pervasive historical introgression alongside ILS during the diversification of the genus *Cervus*.

### 3.4. Non-overlapping window topology weighting and block structure

Across continuous non-overlapping windows (50–1,000 kb), topology weighting and contiguous block-length analyses revealed distinct, scale-dependent patterns of phylogenetic discordance among focal quartets within Cervus and lineages surrounding *E. davidianus* (Figure 5, S7).

**Figure 5.**
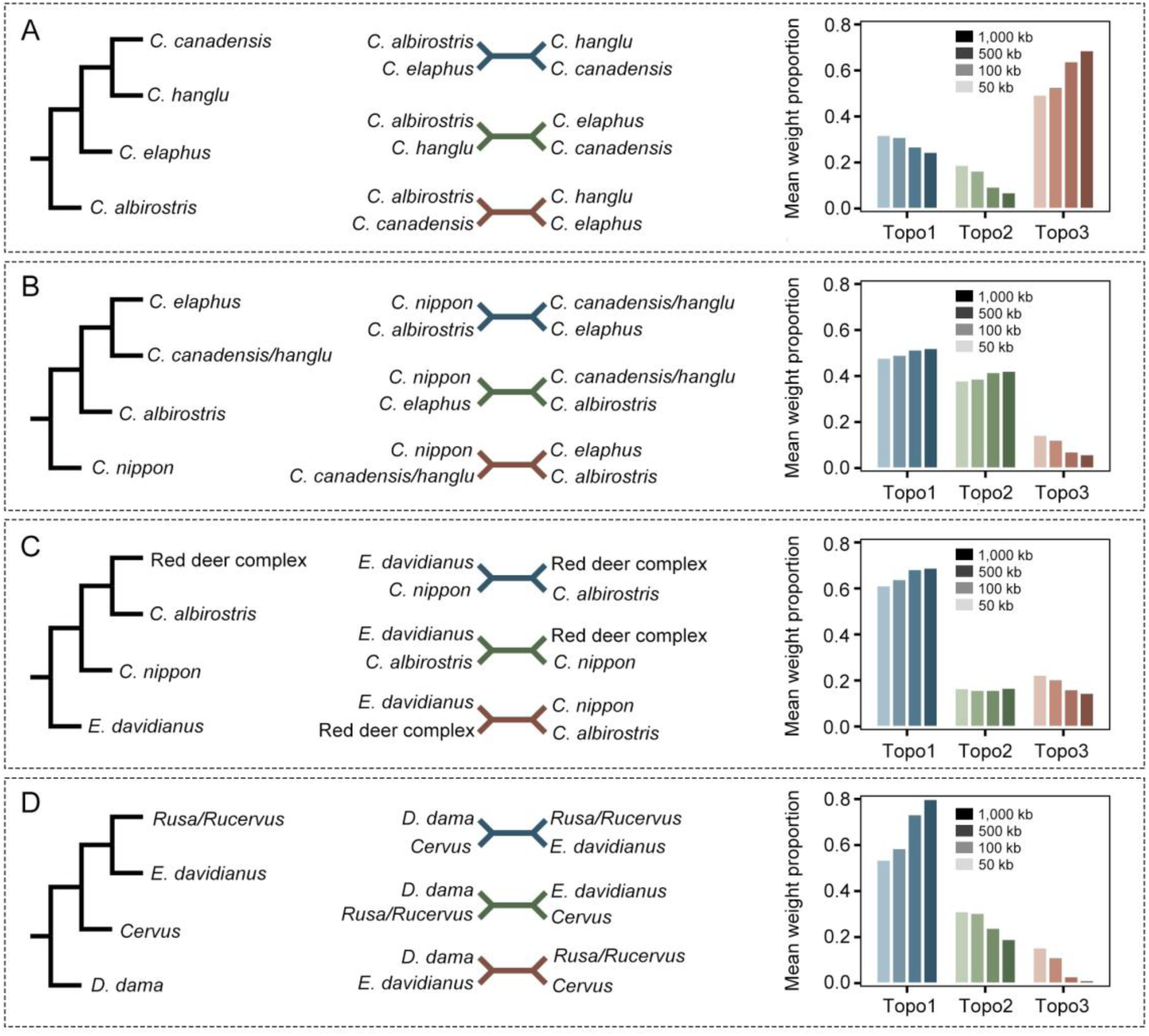
Topology weighting across multiple sliding-window sizes for key nodes in Cervinae. Evaluation of local phylogenetic discordance across different non-overlapping window sizes (50, 100, 500, and 1,000 kb). Panels show quartet evaluations for internal nodes within the genus *Cervus* (A–C) and the lineage subtending *E. davidianus* (D). For each panel, the left diagram illustrates the reference coalescent species tree topology; the middle shows the three possible unrooted quartet topologies (Topo1, Topo2, and Topo3); and the right bar chart depicts the genome-wide mean weight proportions for each topology across the four window sizes. Shaded bars from light to dark represent increasing window sizes from 50 kb to 1,000 kb.

Within the genus *Cervus*, the four-taxon statement examining *C. albirostris* relative to *C. elaphus*, *C. hanglu*, and *C. canadensis* revealed substantial conflict with the species tree (Figure 5A). The alternative discordant topology Topo3 showed a markedly higher genome-wide mean weight than the concordant species-tree topology (Topo1) across all window sizes, with its average weight increasing from ∼0.50 at 50 kb to nearly 0.70 at 1,000 kb. Consistent with this pronounced weighting, genomic blocks supporting Topo3 were exceptionally long and reached multi-megabase spans (up to tens of megabases at larger window sizes), substantially exceeding the fragmented block lengths observed for Topo1 and Topo2 (Figure S7A).

By contrast, the remaining two *Cervus* quartets maintained predominant support for the species tree topology across all window scales (Figure 5B, C). For the node evaluating the placement of *C. nippon* and *C. albirostris* against *C. elaphus* and *C. canadensis*/*C. hanglu*, the concordant topology Topo1 retained the highest mean weight (∼0.50), followed closely by Topo2 (∼0.40), whereas Topo3 contributed minimally (<0.15; Figure 5B). Similarly, in the quartet involving *E. davidianus*, *C. nippon*, *C. albirostris*, and the red deer complex, Topo1 accounted for ∼0.60–0.70 of the mean weight, with Topo2 and Topo3 remaining at low, stable levels (∼0.15–0.20; Figure 5C). In both comparisons, contiguous genomic tracts supporting Topo1 were markedly longer and more extensive than those supporting the alternative topologies, reflecting coherent phylogenetic support across the chromosomes (Figure S7B, C).

For the intergeneric placement of *E. davidianus* relative to *D. dama*, the *Rusa*/*Rucervus* clade, and *Cervus*, the concordant species-tree topology (Topo1) dominated across all window scales, with its mean weight progressively rising from ∼0.55 at 50 kb to ∼0.80 at 1,000 kb (Figure 5D). Concurrently, the weights of the two alternative topologies declined with increasing window size, with Topo2 (*E. davidianus* sister to Cervus) retaining higher proportions than Topo3. Distribution analysis of contiguous segment lengths corroborated this pattern: genomic regions supporting Topo1 formed extensive, continuous chromosomal blocks, whereas regions supporting the two alternative topologies were predominantly confined to short tracts (Figure S7D).

## 4. Discussion

### 4.1. Phylogenomic relationships and taxonomic implications in Cervinae

The generic boundaries and intergeneric relationships within Cervinae have long been debated, partly because morphological classifications based on antler architecture, cranial morphology, and dentition have been complicated by ecological convergence (Brooke, 1878, Groves and Grubb, 1987, Heckeberg, 2020). Subsequent mitochondrial studies further challenged classical taxonomy by revealing widespread paraphyly and conflicting generic placements (Pitra et al., 2004, Hassanin et al., 2012). By analyzing 2,323 sliding-window phylogenetic trees derived from autosomal regions under the multispecies coalescent framework, our study provides a genome-wide species tree framework for Cervinae (Figure 2B). Our results recover *A. porcinus* and *D. dama* as successive early-diverging lineages, with estimated divergence times of 7.69 and 6.56 Ma, respectively, followed by the split between the *Elaphurus*–*Rusa*–*Rucervus* clade and *Cervus* at 5.29 Ma. Their relatively high pairwise genetic distances, exceeding the subfamily-wide mean value of 0.01322 (Figure S3), further reflect the substantial evolutionary divergence of these two lineages. PhyTop analyses also indicate that these deeper divergences are generally well resolved, with discordant topologies showing patterns more consistent with symmetric ILS than asymmetric introgression (Figure 4A).

Our nuclear phylogeny further clarifies the evolutionary relationships and taxonomic positions of several contentious Cervinae lineages. Although *Elaphurus, Rucervus*, and *Rusa* have sometimes been subsumed within a broadly defined *Cervus* (Groves and Grubb, 1987, Hassanin et al., 2012, Hu et al., 2019), our coalescent species tree recovers these three genera as a distinct monophyletic clade that diversified during the Pliocene (4.47–3.36 Ma, Figure 2B), definitively supporting their generic status outside of *Cervus*. Within *Cervus*, our results place *C. albirostris*, formerly assigned to the monotypic genus *Przewalskium* based on its distinctive morphological adaptations to the Qinghai–Tibetan Plateau (Groves and Grubb, 1987), firmly within the genus, diverging at 1.56 Ma after *C. nippon* (2.23 Ma). Its relatively low genetic distances to other *Cervus* species further support this placement (Figure S3). We also recover *C. elaphus*, *C. hanglu*, and *C. canadensis* as distinct evolutionary lineages that diversified around the Early–Middle Pleistocene transition (1.23–1.01 Ma), with *C. hanglu*, represented here by the Tarim red deer formerly classified as *C. e. yarkandensis*, sharing a more recent common ancestor with *C. canadensis* than with *C. elaphus* (Figure 2B) (Hu et al., 2019).

Comparisons with previous nuclear studies highlight the importance of genomic sampling and analytical approaches in resolving relationships among rapidly diverging Cervinae lineages. Early studies based on a limited number of short nuclear gene fragments (e.g., fewer than 3 kb across five nuclear exons and introns) provided insufficient genomic representation to consistently resolve short internal branches (Gilbert et al., 2006, Heckeberg, 2020). Meanwhile, Hu et al. (2019) used reduced-representation genome sequencing (RRGS; approximately 3% genome coverage and 197,543 SNPs) to reconstruct a concatenated ML tree of *Cervus* sensu lato. Their topology differed from our coalescent species tree, particularly in placing *C. albirostris* as the earliest-diverging lineage within *Cervus* sensu stricto and recovering *C. nippon* as sister to the red deer complex. Their divergence-time estimates were also substantially older than ours, potentially reflecting differences in genomic sampling, phylogenetic reconstruction, and molecular dating calibrations. Topological differences were also evident between our coalescent species tree, the NJ tree (Figure S2), and the mitochondrial ML tree (Figure 2C). Unlike the multispecies coalescent, which explicitly accommodates gene-tree discordance arising from ILS, NJ and concatenated ML approaches do not model heterogeneous genealogical histories across genomic regions. Consequently, their inferred topologies may be influenced by the combined effects of ancestral polymorphism and historical introgression. In particular, allele sharing between *E. davidianus* and different *Cervus* species (Figure S6) may contribute to reduced genetic distances and the basal placement of *E. davidianus* in the NJ tree, although this relationship cannot be attributed to introgression alone. Similarly, the mitochondrial genome represents a single maternally inherited, non-recombining locus and may retain a distinct evolutionary history from the nuclear genome, particularly following mitochondrial capture through hybridization (Pitra et al., 2004, Heckeberg, 2020). Together, these comparisons support the use of genome-wide data and coalescent-based approaches to establish a reliable species-tree framework, while highlighting the need to consider historical gene flow separately when interpreting the complex evolutionary history of Cervinae.

### 4.2. Intergeneric phylogenetic discordance and historical gene flow in Cervinae

The early intergeneric diversification of Cervinae appears to have been primarily affected by extensive ILS, whereas later divergences may have involved both ancestral polymorphism and post-divergence gene flow. Our Phytop analyses reveal considerable phylogenetic discordance across deep intergeneric nodes. At the earliest divergences, such as those involving *A. porcinus* and *D. dama*, the three alternative topologies occurred at relatively similar frequencies, approaching the expected 1/3 under extensive ancestral ILS. Although PhyTop estimated relatively high introgression indices (IH-i) at some of these nodes, including 38.8% at the divergence of *D. dama*, such estimates should be interpreted with caution. Under conditions of extensive ancestral ILS, when the dominant topology has relatively low support, small differences in alternative topology frequencies can produce apparent introgression signals (Shang et al., 2024). Thus, the relatively symmetric distribution of alternative topologies at these early nodes is more consistent with extensive ancestral ILS than with substantial asymmetric gene flow (Figure 4A). In contrast, the divergence involving *E. davidianus* and the *Rusa– Rucervus* lineage shows evidence of both ILS (ILS-i = 47.2%) and introgression (IH-i = 27.8%). This suggests that, in addition to retaining ancestral polymorphism, the lineage leading to *E. davidianus* may have experienced gene flow with *Cervus* species or its ancestral populations after its divergence. Such gene flow may have contributed to the varying phylogenetic placement of *E. davidianus* and *R. alfredi* among different phylogenetic analyses (Figure 2B, 2C, S2).

The possible role of historical gene flow in the evolution of *E. davidianus* is particularly relevant to the long-standing hypothesis that this species originated through hybridization between *R. eldii* and *C. canadensis*, as proposed by Meijaard and Groves (2004) and Pitra et al. (2004). Our ABBA-BABA tests revealed excess allele sharing between *E. davidianus* and *R. eldii* (Figure S4, Table Sx). However, the similar D statistics obtained when each of the five sampled *Cervus* species was used as H3 suggest that this signal is not specifically associated with any one extant *Cervus* species (Figure S6). With *R. eldii* as H1, D values ranged from 0.1381 to 0.1428, while those with *R. alfredi* as H1 ranged from 0.0928 to 0.1148 (Table Sx). Consistent with this pattern, *f*-branch analysis also showed relatively similar *f*_b_ values between *E. davidianus* and the five sampled *Cervus* species. Together, these results suggest that the gene flow involving *E. davidianus* may have originated from an ancestral *Cervus* population rather than from a specific extant *Cervus* species. Therefore, although historical gene flow likely contributed to the evolutionary history of *E. davidianus*, our results do not provide direct support for the specific hypothesis that it originated through hybridization between *R. eldii* and a particular *Cervus* species.

### 4.3. Incomplete lineage sorting and recent introgression shape phylogenetic relationships within *Cervus*

Although our dataset includes only one representative species for basal genera such as *Axis* and *Dama*, the sampling of five species within *Cervus* allows us to investigate intrageneric diversification in greater detail. These five lineages diverged within a relatively short period of approximately 1.2 million years during the Early Pleistocene (2.23–1.01 Ma, Figure 2B). Such rapid diversification likely resulted in the retention of substantial ancestral polymorphism, leading to high levels of ILS across internal nodes (ILS-i = 38.9%–51.9%, Figure 4A). At the basal divergence of *Cervus*, PhyTop detected no evidence of asymmetric introgression from the *E. davidianus–Rusa–Rucervus* lineage (IH-i = 0.0%, Figure 4A). Although D-statistics showed some variation in allele-sharing patterns when *E. davidianus* was used as H3 (D = 0.0076–0.0682, Figure S6, Table Sx), these signals do not necessarily indicate direct gene flow between *E. davidianus* and a particular *Cervus* species. Instead, they may reflect the combined effects of ILS and historical gene flow among *Cervus* lineages. Within *Cervus*, however, PhyTop detected substantial introgression signals in addition to extensive ILS (IH-i = 36.7%–44.7%, Figure 4A), suggesting that post-divergence gene flow played an important role in shaping the genomic relationships among species.

Our multi-scale sliding-window topology-weighting analyses and contiguous block-length comparisons provide further insights into the genomic distribution of these discordant relationships (Figure 5, S7). In particular, the changes in topology weights with increasing window size, together with the lengths of consecutive genomic blocks supporting alternative topologies, help distinguish the genomic patterns associated with ancestral ILS and more recent introgression. Ancestral polymorphisms retained through successive speciation events are expected to be fragmented by recombination over time, whereas relatively recent introgression can preserve longer genomic segments carrying the ancestry of another lineage. Consequently, introgression-related topologies may receive greater support when larger genomic windows encompass these long, linked segments. This pattern is particularly evident in the quartet comprising *C. albirostris*, *C. elaphus*, *C. hanglu*, and *C. canadensis* (Figure 5A). Rather than the species-tree topology (Topo1), the alternative unrooted topology (Topo3), which groups *C. albirostris* with *C. canadensis* and *C. hanglu* with *C. elaphus*, became increasingly dominant as window size increased. Its mean weight rose from approximately 0.50 at 50 kb to nearly 0.70 at 1,000 kb, whereas the weight of Topo1 declined over the same range. Consistently, genomic blocks supporting Topo3 were substantially longer than those supporting Topo1, extending across multi-megabase regions (Figure S7A). These patterns suggest that the alternative topology is supported by relatively long genomic segments, consistent with substantial historical introgression. Because this unrooted topology can correspond to two rooted relationships, namely ((*C. albirostris*, *C. canadensis*), (*C. hanglu*, *C. elaphus*)) and (*C. albirostris*, (*C. canadensis*, (*C. hanglu*, *C. elaphus*))), its increased weight may reflect allele sharing involving more than one pair of lineages. This interpretation is consistent with the strong allele-sharing signal between *C. albirostris* and *C. canadensis* (*f_b_* > 0.2; maximum D = 0.3266), as well as evidence of gene flow involving *C. hanglu* and *C. elaphus* (Figure 3, S4). A similar increase in the weight of the alternative topology grouping *C. albirostris* with the *C. canadensis/C. hanglu* lineage was observed in the quartet involving *C. nippon* and the red deer complex (Figure 5B). Together, these results suggest that relatively long introgressed genomic segments have contributed to the increased representation of alternative topologies at larger window sizes.

Beyond the strong allele-sharing signal between *C. albirostris* and *C. canadensis*, our D-statistics and *f*-branch analyses reveal a broader network of historical gene flow within *Cervus* (Figure 3, S4, S5). Both *C. albirostris* and *C. nippon* show excess allele sharing with *C. canadensis*, as well as with each other. These interconnected signals may have contributed to the mitochondrial placement of *C. albirostris* and *C. nippon* within the *C. canadensis* clade (Figure 2C). However, because gene flow was also detected between *C. albirostris* and *C. nippon*, the shared signals among these species may have resulted from direct or indirect gene exchange, and their respective contributions to the mitochondrial relationships remain unresolved. Furthermore, although D-statistics indicate greater allele sharing of both *C. albirostris* and *C. nippon* with *C. hanglu* than with *C. elaphus*, *f*-branch analysis places this signal on the ancestral branch shared by *C. hanglu* and *C. canadensis* (Figure 3). This suggests that excess allele sharing may be associated with gene flow involving their ancestral population, rather than necessarily reflecting direct introgression from extant *C. hanglu*. Overall, our results indicate that the genomic evolution of *Cervus* was shaped by rapid lineage diversification, extensive ancestral ILS, and subsequent gene flow among diverging lineages. The resulting mosaic of genealogical histories helps explain the substantial differences among phylogenetic relationships inferred from different genomic regions.

### 4.4. Climatic changes and the reticulate evolutionary history of Cervinae

The divergence time estimates provide a temporal framework for examining the relationship between Cervinae diversification and past environmental changes. Our divergence time estimates provide a stable temporal framework for reconstructing the evolutionary history of Cervinae, as the major divergence times are broadly consistent with those reported by Sinding et al. (2026) based on a subset of cervid species and a ruminant study (Chen et al., 2019). Within this framework, the diversification of Cervinae extends from the Late Miocene to the Pleistocene, with successive divergences of basal lineages followed by rapid diversification within *Cervus*. These events broadly coincided with major climatic and ecological transitions across Eurasia. Specifically, the initial divergence of Cervinae overlapped with Late Miocene global cooling, intensified mid-latitude aridification, and the expansion of open habitats across Eurasia (11–7 Ma) (Eronen et al., 2009, Wang et al., 2025). Because cervids are predominantly brachydont herbivores adapted to forest edges and mesic environments, these arid open biomes, which favored hypsodont ungulates such as bovids and equids, constrained early cervid radiation, resulting in the emergence of only basal lineages such as *Axis* (7.69 Ma) and *Dama* (6.56 Ma) during this interval. By contrast, the subsequent divergence between the *Elaphurus*–*Rusa*–*Rucervus* clade and *Cervus* occurred near the Miocene–Pliocene boundary (5.29 Ma), followed by the internal radiation of *Elaphurus*, *Rusa*, and *Rucervus* during the Pliocene (4.47–3.36 Ma; Figure 2B), as well as the divergence between *Dama* and extinct *Megaloceros* (∼3.5 Ma) (Sinding et al., 2026). This timing coincides with a climatic reversal toward increased humidity and forest expansion around ∼5 Ma (Eronen et al., 2009), which likely released ecological niches for forest– and wetland-adapted deer.

Together, our results suggest that the evolutionary history of Cervinae was shaped by the interplay between successive lineage diversification, extensive ancestral ILS, and recent introgression. Cladogenesis over short evolutionary intervals generated a high level of ILS, while subsequent climatic and geographic changes may have repeatedly brought diverged lineages into secondary contact, allowing interspecific gene flow. The resulting introgressed genomic segments, particularly those retained as relatively long contiguous blocks, contributed to the heterogeneous distribution of genealogical relationships across the genome. This combination of persistent ancestral polymorphism and episodic gene flow produced a reticulate evolutionary history that cannot be fully represented by a strictly bifurcating species tree. Our findings therefore highlight how continental radiation and recent secondary contact can jointly shape genomic variation and phylogenetic relationships in diversifying mammalian groups.

## Author Contributions

Conceptualization, B. X., G. S.; Data collection, B. X., X. S.; Data analyses, B. X.; Computing platform, G. S., J. Y.; Writing-Original Draft, B. X., G. S.; Writing-Review and Editing, All authors; Supervision, G. S.; Funding Acquisition, G. S., J. Y..

## Acknowledgements

Guilian Sheng and Junxia Yuan were funded by the National Natural Science Foundation of China (Grant 42572005 and 42472008).

## Data Availability Statement

All accession numbers of published data used in this manuscript were provided in the Supporting Information.

