## Supplemental Figures and Table S1 for "Shifting roles of incomplete lineage sorting and gene flow across the diversification of Old World deer"

**This file includes the following:**

**Figure S1**. Sensitivity analysis of divergence time estimates to the low-coverage *R. eldii* genome.

**Figure S2**. NJ phylogenetic tree of Cervinae based on LD-pruned autosomal SNPs.

**Figure S3**. Genome-wide pairwise identity-by-state (IBS) genetic distances among Cervinae species.

**Figure S4**. Summary of D-statistics across all evaluated taxon combinations in Cervinae.

**Figure S5**. Genome-wide D-statistics evaluating excess allele sharing among Cervus species.

**Figure S6**. Genome-wide D-statistics evaluating excess allele sharing among *E. davidianus*, *R. eldii*, *R. alfredi*, and *Cervus* species.

**Figure S7**. Distribution of contiguous genomic segment lengths supporting alternative topologies across multiple window sizes.

**Table S1**. Public sequence data used in this study.

**Table S2**. MCMCtree divergence times of all Cervinae splits (Ma).

**Table S3**. P-distance among all Cervinae species pairs.

**Table S4**. D statistics of all Cervinae species combinations.

**SM References**

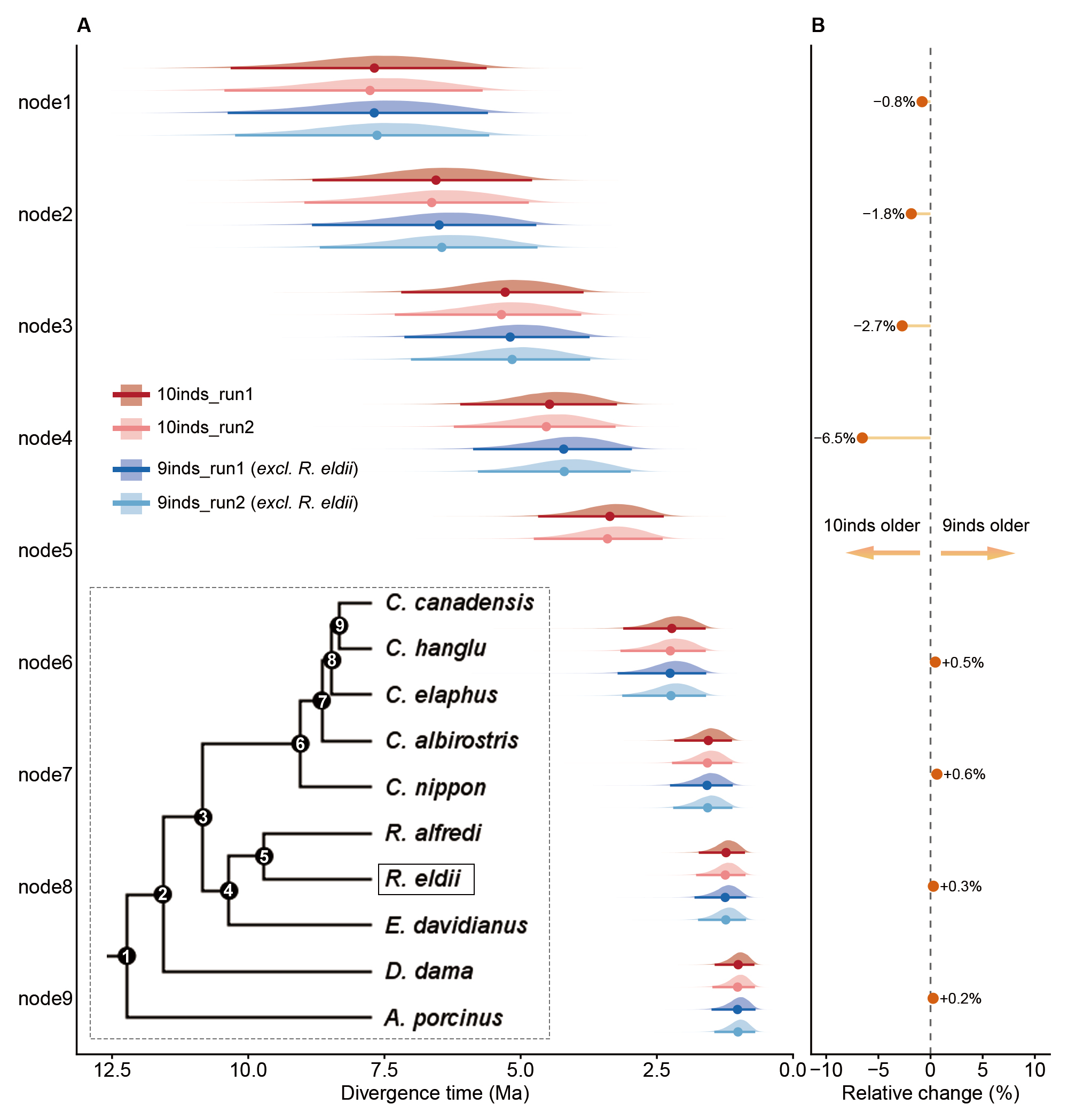

**Figure S1**. Sensitivity analysis of divergence time estimates to the low-coverage *R. eldii* genome. (A) Posterior distributions of divergence times across internal nodes of Cervinae estimated from two independent MCMCtree runs for the full 10-species dataset (10inds_run1, 10inds_run2; red) and the reduced 9-species dataset excluding *R. eldii* (9inds_run1, 9inds_run2; blue). Dots and horizontal bars represent posterior means and 95% highest posterior density intervals, respectively. The inset cladogram indicates the positions of the numbered internal nodes (node1–node9), with the boxed taxon (*R. eldii*) excluded in the 9-species analysis. (B) Relative percentage changes in posterior mean divergence times across the eight shared nodes after excluding *R. eldii*. Positive and negative values indicate older and younger age estimates in the 9-species dataset, respectively.

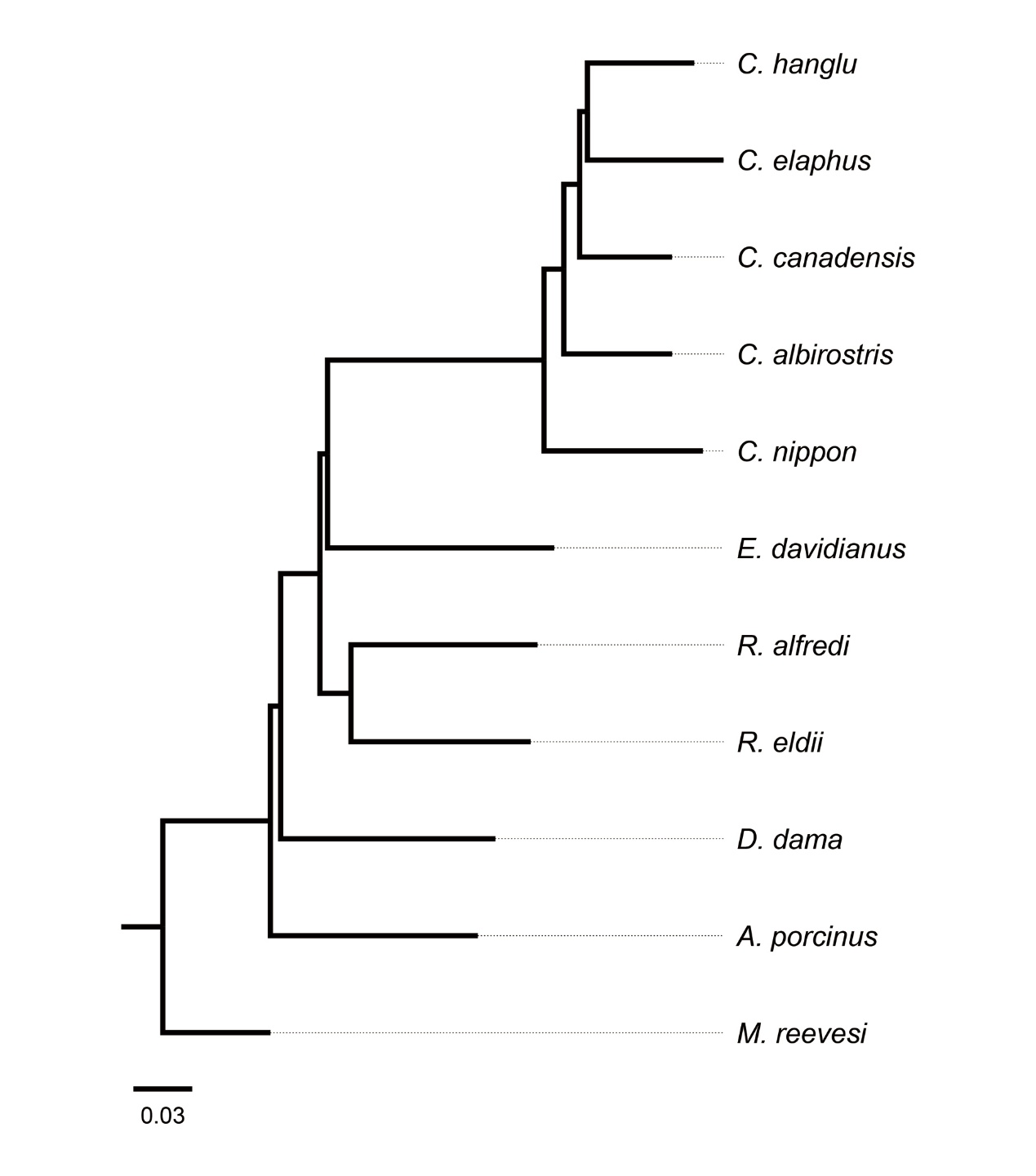

**Figure S2**. NJ phylogenetic tree of Cervinae based on LD-pruned autosomal SNPs. The tree was constructed from pairwise genetic distances calculated across 10,041,723 LD-pruned SNPs (r*^2^* < 0.2$), rooted with *M. reevesi* as the outgroup. The scale bar indicates genetic distance (0.03 substitutions per site).

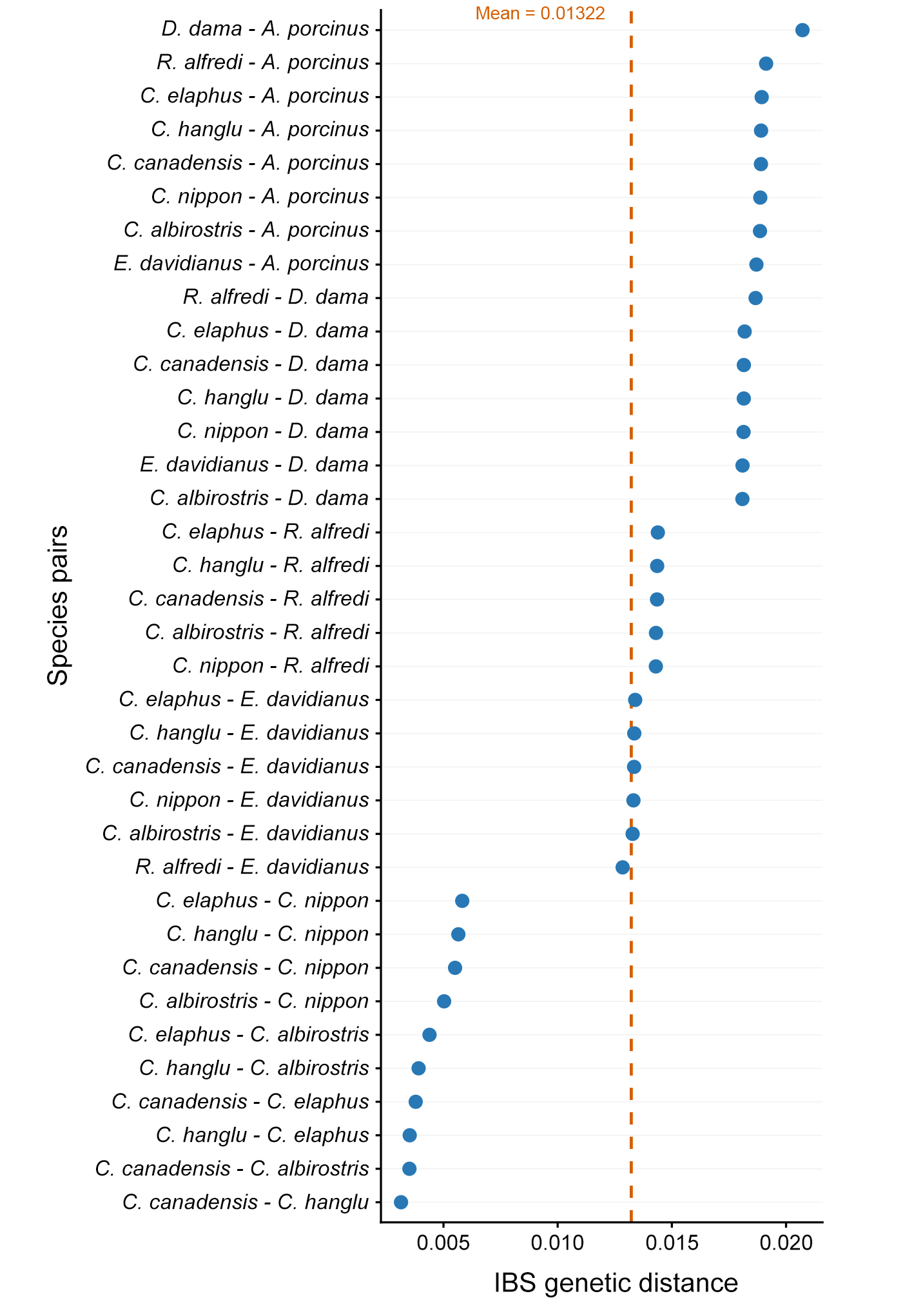

**Figure S3**. Genome-wide pairwise identity-by-state (IBS) genetic distances among Cervinae species. Blue dots represent pairwise autosomal IBS distances calculated across all 36 species pairs among nine Cervinae species (excluding the low-coverage *R. eldii* genome). The vertical orange dashed line indicates the overall mean genetic distance across all pairs (Mean = 0.01322)

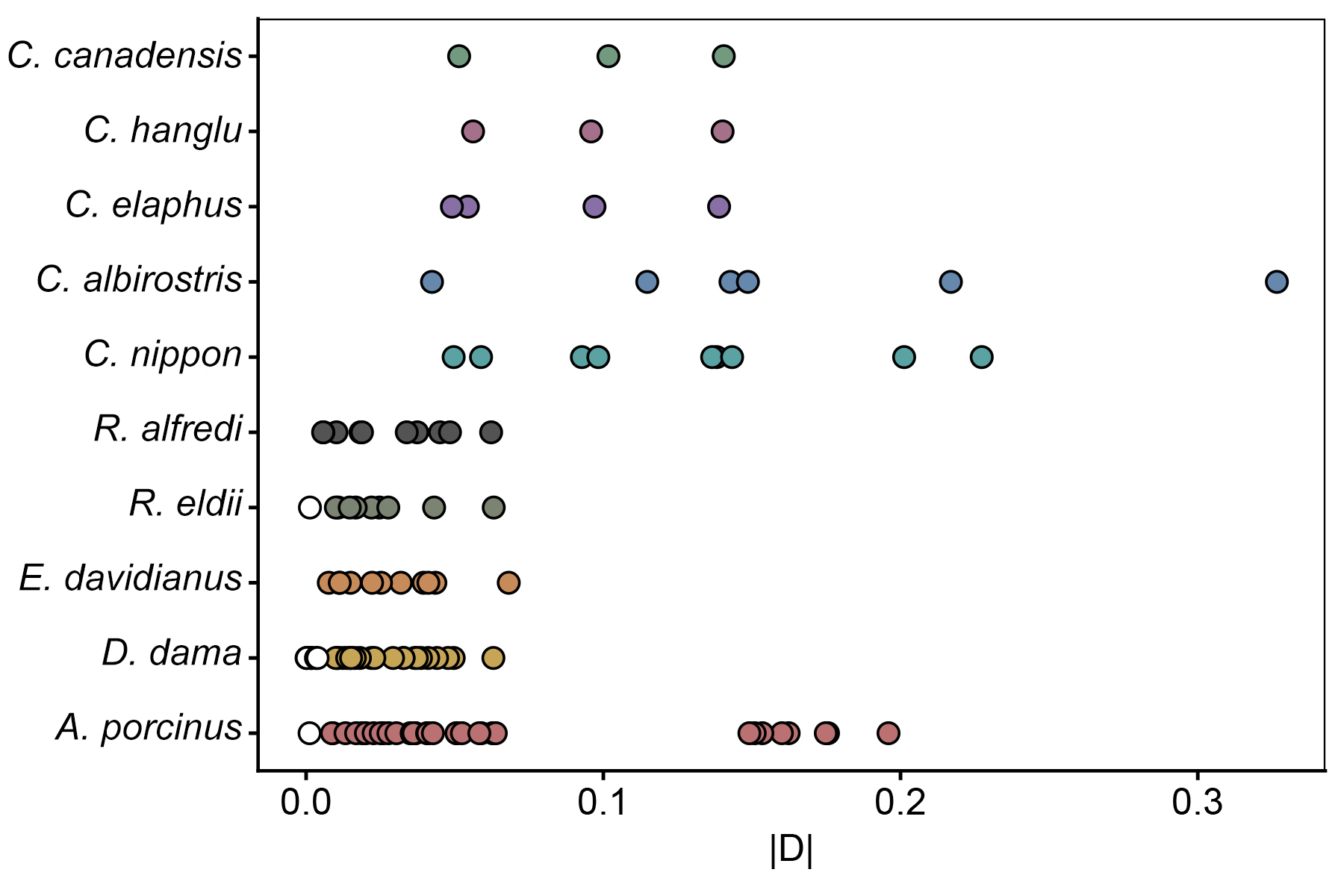

**Figure S4**. Summary of D-statistics across all evaluated taxon combinations in Cervinae. Each row shows the distribution of absolute D values (|D|) for all tested three-taxon combinations (as ((P1, P2), P3), *M. reevesi*) associated with each focal species. Filled colored circles indicate statistically significant introgression signals (|Z| ≥ 3; 113 of 121 combinations), whereas open circles denote non-significant tests (|Z| < 3).

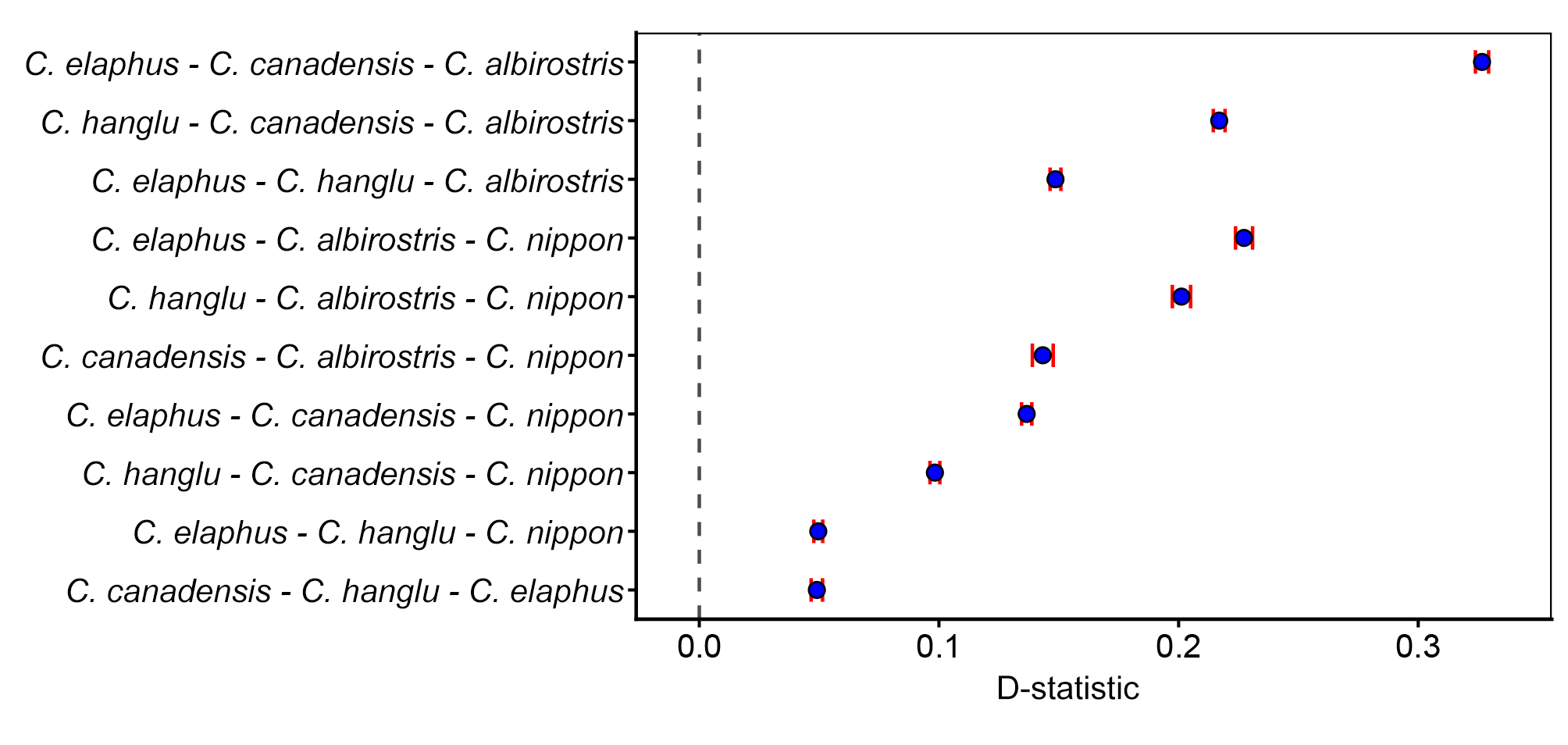

**Figure S5**. Genome-wide D-statistics evaluating excess allele sharing among Cervus species. Blue dots represent D-statistic estimates for selected three-taxon combinations arranged along the y-axis as P1 – P2 – P3, with *M. reevesi* serving as the outgroup (P4). Horizontal red error bars indicate standard errors estimated via block jackknifing across 1-Mb genomic blocks, and the vertical dashed line denotes D = 0. Positive D values (D = 0.0496–0.3266; all |Z| ≥ 3) indicate significant excess allele sharing between P2 and P3 relative to P1.

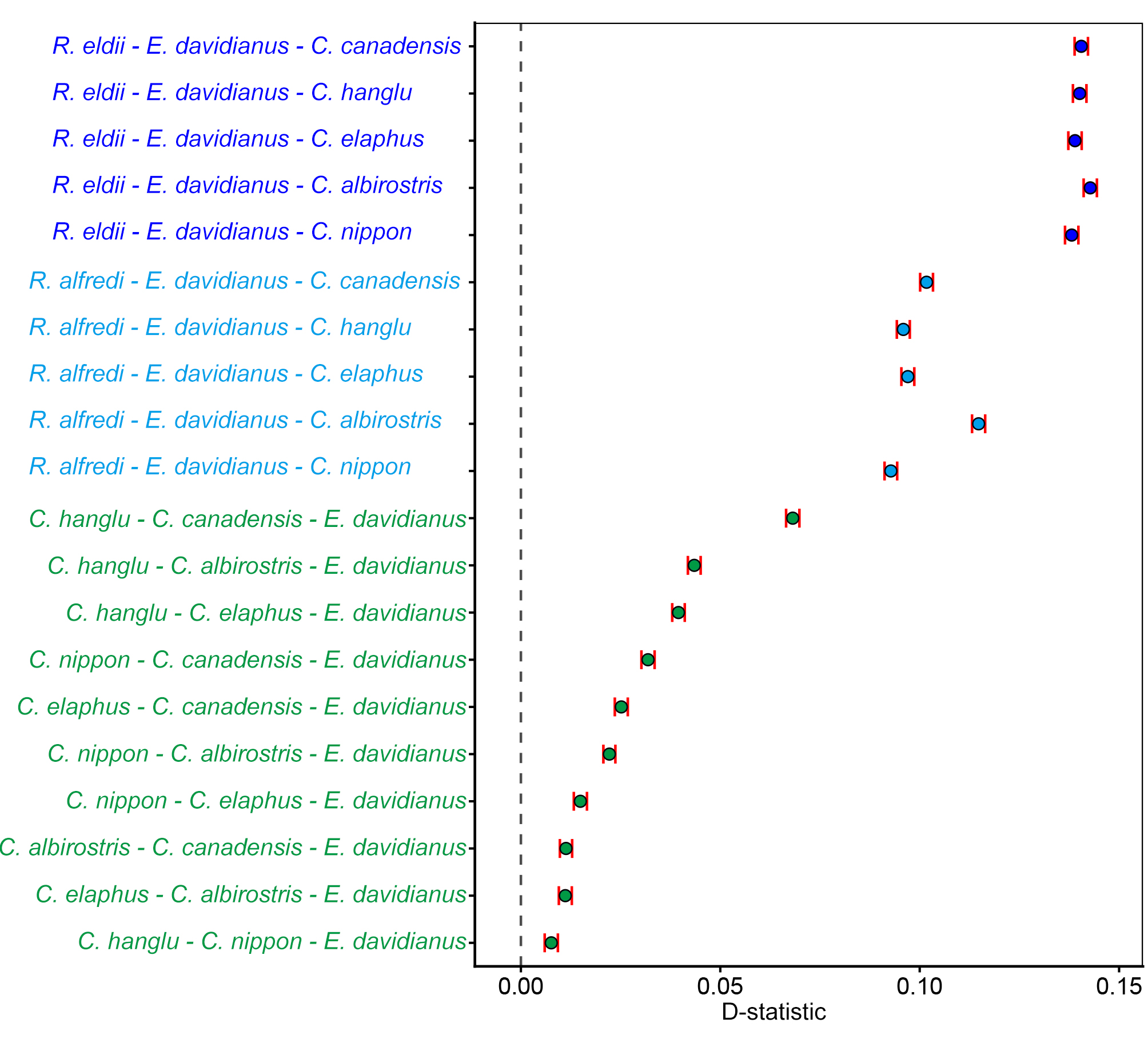

**Figure S6**. Genome-wide D-statistics evaluating excess allele sharing among *E. davidianus*, *R. eldii*, *R. alfredi*, and *Cervus* species. Blue dots represent D-statistic estimates for selected three-taxon combinations arranged along the y-axis as P1 – P2 – P3, with *M. reevesi* serving as the outgroup (P4). Horizontal red error bars indicate standard errors estimated via block jackknifing across 1-Mb genomic blocks, and the vertical dashed line denotes D = 0. Positive D values (all |Z| ≥ 3) indicate significant excess allele sharing between P2 and P3 relative to P1.

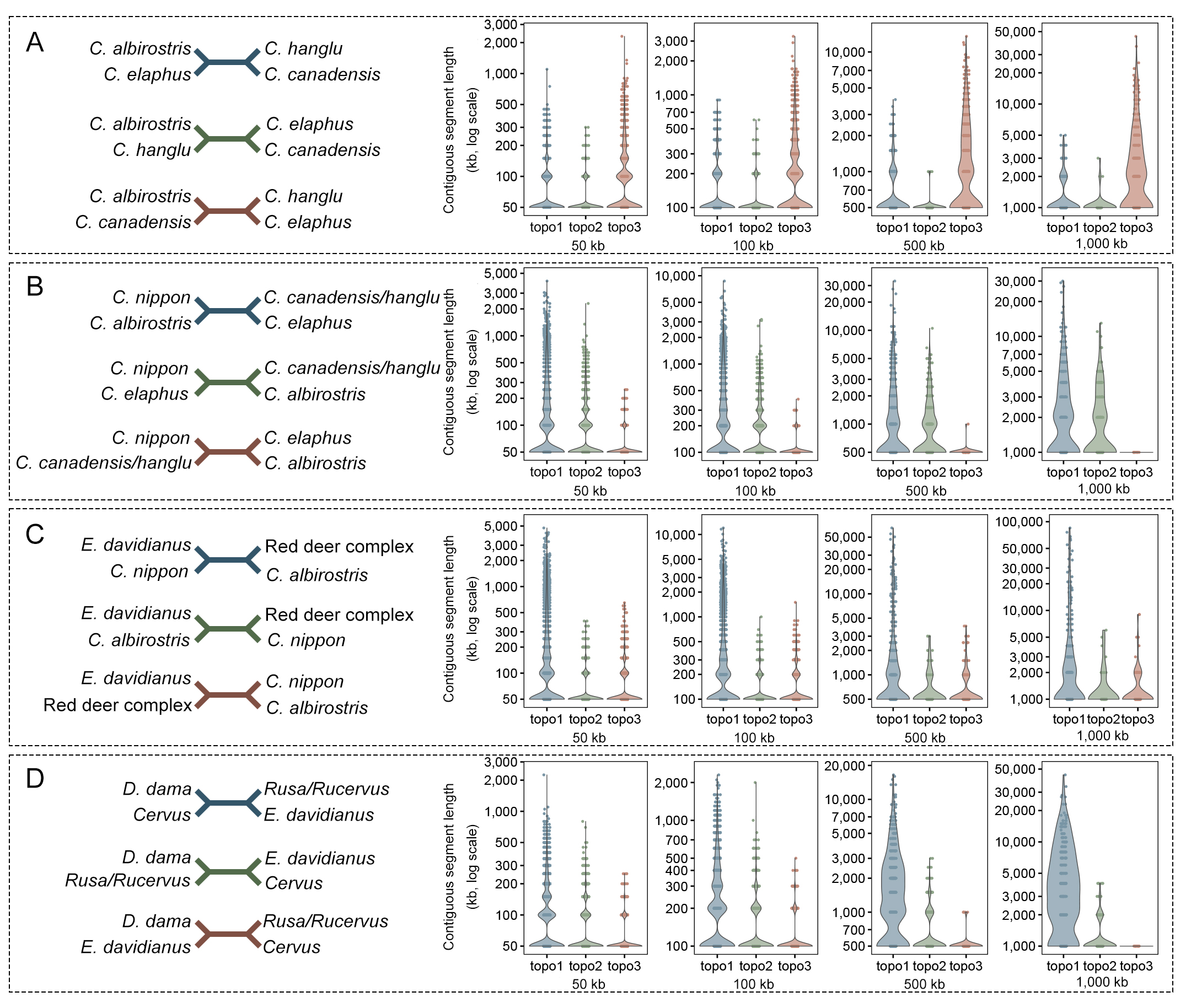

**Figure S7**. Distribution of contiguous genomic segment lengths supporting alternative topologies across multiple window sizes. Analysis of contiguous block lengths supporting each of the three unrooted topologies (topo1, topo2, and topo3) evaluated across four non-overlapping sliding-window scales (50, 100, 500, and 1,000 kb). (A–C) Quartet evaluations for internal nodes within the genus *Cervus*. (D) Quartet evaluation for the lineage subtending *E. davidianus*. For each panel, the left diagram depicts the corresponding unrooted topologies; the violin and jitter plots on the right display the distributions of contiguous fragment lengths (in kb, log10 scale) supporting each topology under increasing window sizes (from left to right: 50, 100, 500, and 1,000 kb). Horizontal lines within violins indicate the median and interquartile ranges.

**Table S1**. Public sequence data used in this study.

| Species | Accession No. | Mean depth | Sex | Reference |
| --- | --- | --- | --- | --- |
| *C. hanglu* | SAMN21971318 | 28.1289 | F | Bergeron et al. (2023) |
| *C. elaphus* | SAMN44812736 | 21.0533 | F | de Jong et al. (2025) |
| *C. canadensis* | SAMC4592118 | 28.0785 | M | Zhong et al. (2025) |
| *C. albirostris* | SAMN08714477 | 17.1855 | M | Chen et al. (2019) |
| *C. nippon* | SAMN21971320 | 11.2101 | M | Bergeron et al. (2023) |
| *E. davidianus* | SAMN31632925 | 17.2577 | M | Chen et al. (2019) |
| *R. alfredi* | SAMN41012784 | 22.8701 | M | Direct submission |
| *R. eldii* | SAMC1224815 | 2.7807 | F | Zheng et al. (2024) |
| *D. dama* | SAMN31886425 | 22.818 | M | Barnard et al. (2023) |
| *A. porcinus* | SAMC4592122 | 32.0163 | M | Zhong et al. (2025) |
| *M. reevesi* | SAMN11611988 | 17.4073 | M | Mudd et al. (2020) |

**Table S2**. MCMCtree divergence times of all Cervinae splits (Ma).

*Note: provided in a separate Excel file.*

**Table S3**. P-distance among all Cervinae species pairs.

*Note: provided in a separate Excel file.*

**Table S4**. D statistics of all Cervinae species combinations.

*Note: provided in a separate Excel file.*
